# Mapping the growth effect of previously hidden ubiquitin alleles using an overexpression based mutational scan

**DOI:** 10.1101/831503

**Authors:** David Mavor, Daniel N. A. Bolon, Parul Mishra

## Abstract

Deep mutational scanning has emerged as a powerful, high throughput approach to determine the growth effect of thousands of alleles at once in bulk competition. However, to date only the growth effect of mutant alleles in isolation has been determined. Building off previous work, we have created a library of all possible single point mutations in ubiquitin and determined the growth effect of mutants overexpressed in the presence of a wild type allele. Using this scan, we explained over half of the previously missing mutants in the single allele scan by showing that they exhibit deleterious effects when co-expressed with wild type ubiquitin. Additionally, unlike the single allele growth effect, these overexpression growth effects were distributed across the entire protein. This overexpression scan methodology can identify likely dominant mutant effects in any essential gene and is highly complementary with traditional deep mutational scanning approaches.

## Introduction

In order to interrogate the sequence-function relationships of proteins, investigators have increasingly turned to high throughput, deep sequencing based mutational scans. Termed Deep Mutational Scanning (DMS), these studies have elucidated the functional impact of individual point mutations via either a direct functional readout(1) or more commonly by using growth rate as a functional proxy for essential genes(2, 3, 4, 5, 6, 7). While generating point mutant libraries has been possible for years(8), DMS offers a systematic approach quickly to determine the functional consequence of thousands alleles in parallel opening up exciting opeterninites to study genetic interactions(9) and chemical genetics(10, 11).

In previous work we developed an approach to measure the fitness effect of all possible single point mutants of a gene which we term EMPIRC(3, 4). We generate systematic site saturation libraries through the incorporation of a single degenerate codon (NNN) into a wild type (WT) coding sequence. Due to this systematic approach, all possible single point mutants are included in the initial library and the vast majority of the alleles can be observed by deep sequencing. We transform these libraries into a strain harbouring a conditional WT allele that can be strictly regulated. This allows for the library to be expanded under permissive conditions and then the fitness effect of the mutant allele to be determined in bulk competition when the expression of the WT allele is turned off. However, we observed that some alleles which were successfully generated and confirmed where never observed after the initial library outgrowth in permissive conditions. Hypothesising that these mutants were toxic even in the presence of a WT allele, we sought to determine the fitness effect of these missing alleles. Building off of our EMPIRIC study of ubiquitin(12) we inverted the regulatory scheme to determine the fitness effect of all possible single point mutants of ubiquitin in the presence of a WT allele. The library is initially grown with only the WT allele expressed. Bulk competition is started by the overexpression of the mutant alleles and the allele frequencies followed to determine the fitness effect of the mutant in the presence of the WT allele. Here we present the first determination of dominant point mutant growth effects by DMS.

Ubiqutin is an essential eukaryotic protein that is central to protein homeostasis via its ability to be covalently linked to other proteins(13). The C-terminus of ubiquitin is ligated to amine groups of target proteins through the three enzyme cascade of E1, E2, and E3(14, 15). Ubiquitin can also be removed from substrate proteins by deubiquitinating enzymes (DUBs) increasing the complexity of the regulation(16). Additionally, multiple ubiquitin molecules are often ligated together between the C-terminus of one and an amine from another. While any of the seven lysine residues or the N-terminus can be linked and all possible linkages have been observed in yeast(17), only Lysine 48 (K48) appears to be essential under normal growth conditions(18). K48 linked poly-ubiquitin directs substrate proteins to the proteasome for degradation(19, 20) explaining the essential role of ubiquitin in proteostasis which can lead to disease when miss-regulated(21, 22). Of the remaining poly-ubiquitin linkages, only the role of K63 linkages in the DNA damage response(23) and endocytosis(24) are well established in yeast.

## Results

### Inverting selection on the Ub library to determine dominant negative mutations

Previous DMS studies of Ub determined the growth rates of all possible Ub point mutants in yeast grown in rich media(12) or under chemical stress conditions(10, 11). In these selections, each library was grown for multiple generations in the presence of both a mutant allele and the WT allele. In each case, after this outgrowth many alleles were either depleted or missing in the initial sample. We hypothesised that these depleted alleles represent dominant negative or ‘dominant sick’ alleles. Because of this phenotype, the fitnesses of these alleles could not be determined through this single allele scan.

In order to assess the fitness effects of these putative ‘dominant sick’ alleles, the selection method of the library was inverted. Rather than constitutively expressing the WT allele and expressing the mutant under a galactose regulatable promoter, the mutant is constitutively expressed and the WT expression regulated. Therefore when the library is grown in glucose only the WT allele is expressed. Upon transfer of the library to galactose the mutant allele is overexpressed. The bulk competition of the library is performed in galactose with both alleles expressed and the growth rate of each mutant is determined.

### Many mutants display a synthetic sick phenotype when co-expressed with WT Ub

In order to assess the growth rates of all Ub point mutants when overexpressed alongside the WT allele nine samples were taken over 24 hours of selection and the mutant plasmids isolated and prepared for deep sequencing. For each sample,WT normalized mutant allele frequencies were determined and then a growth effect for each allele was calculated. The resulting mutant overexpression fitness landscape shows a striking variability in mutant growth rates (Figure 1).

**Figure 1.**
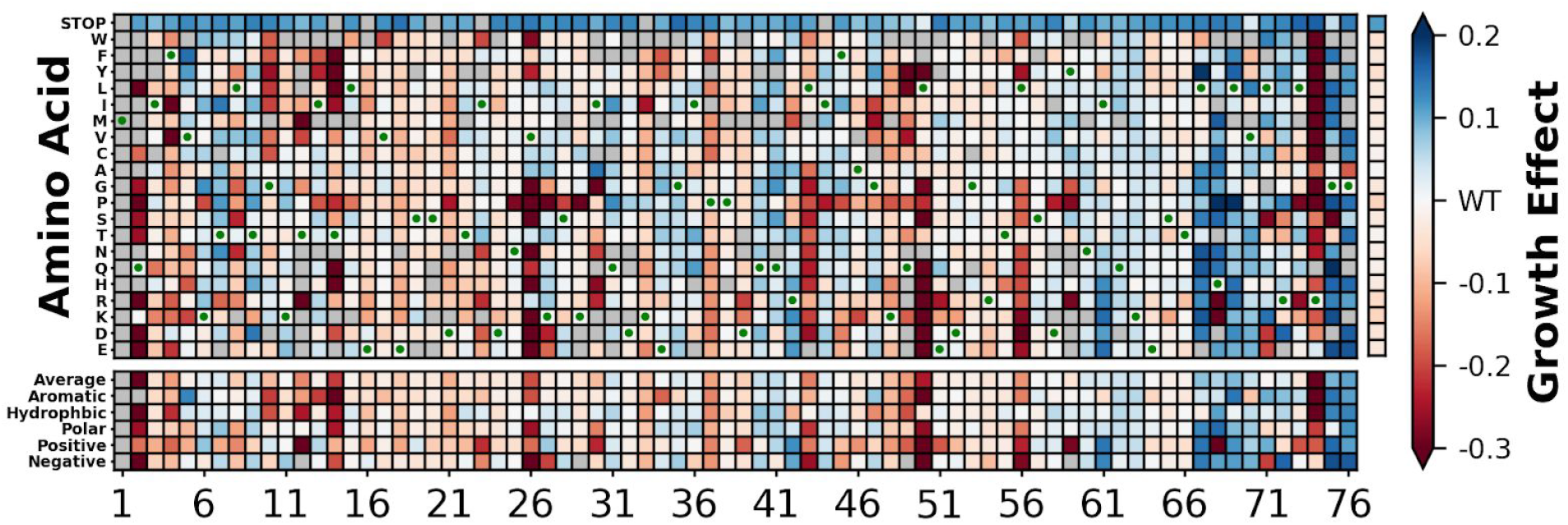
Overexpression of mutant alleles in the presence of WT ubiquitin leads to a variety of growth effects. Growth effects were calculated for each observed allele and presented as a heatmap.WT amino acids are shown with a green dot. Unobserved alleles are shown in grey. Overexpression of the mutants leads to beneficial growth effects (blue) and deleterious growth effects (red) with strong common deleterious effects at positions 2, 14, 26, 43, 50, 56, and 74

Interestingly, many alleles that were deleterious in the single allele scan scan, such as stop codons and essential c-terminal residues, grew faster than WT (Figure 1 - blue boxes). However mutants of the essential Lys48 residue were generally deleterious. Additionally, mutants at several positions (Positions 2, 14, 26, 43, 50, 56, 74) show strong deleterious phenotypes for many mutations (Figure 1 - red boxes).

Additionally, we examined the distribution of the growth effects. This distribution shows a large peak around the WT growth rate with a shoulder around 0.1 representing the stop codon population. Sick alleles are present at growth rates less than around −0.05 followed by a tail down to growth rates around −0.5. Using the distribution of growth effects of the WT synonyms a Z score was calculated for each mutant. Based on a two-tailed Z-test the mutants were binned based on cutoffs of Z > 4 and Z < −4 (Figure 2A). The mutant growth rates were binned into deleterious alleles (growth effect less than WT - red - 37.7%), WT like alleles (growth effects indistinguishable from WT - green - 34.5%), and beneficial alleles (growth rates greater than WT - blue - 27.8%).

**Figure 2.**
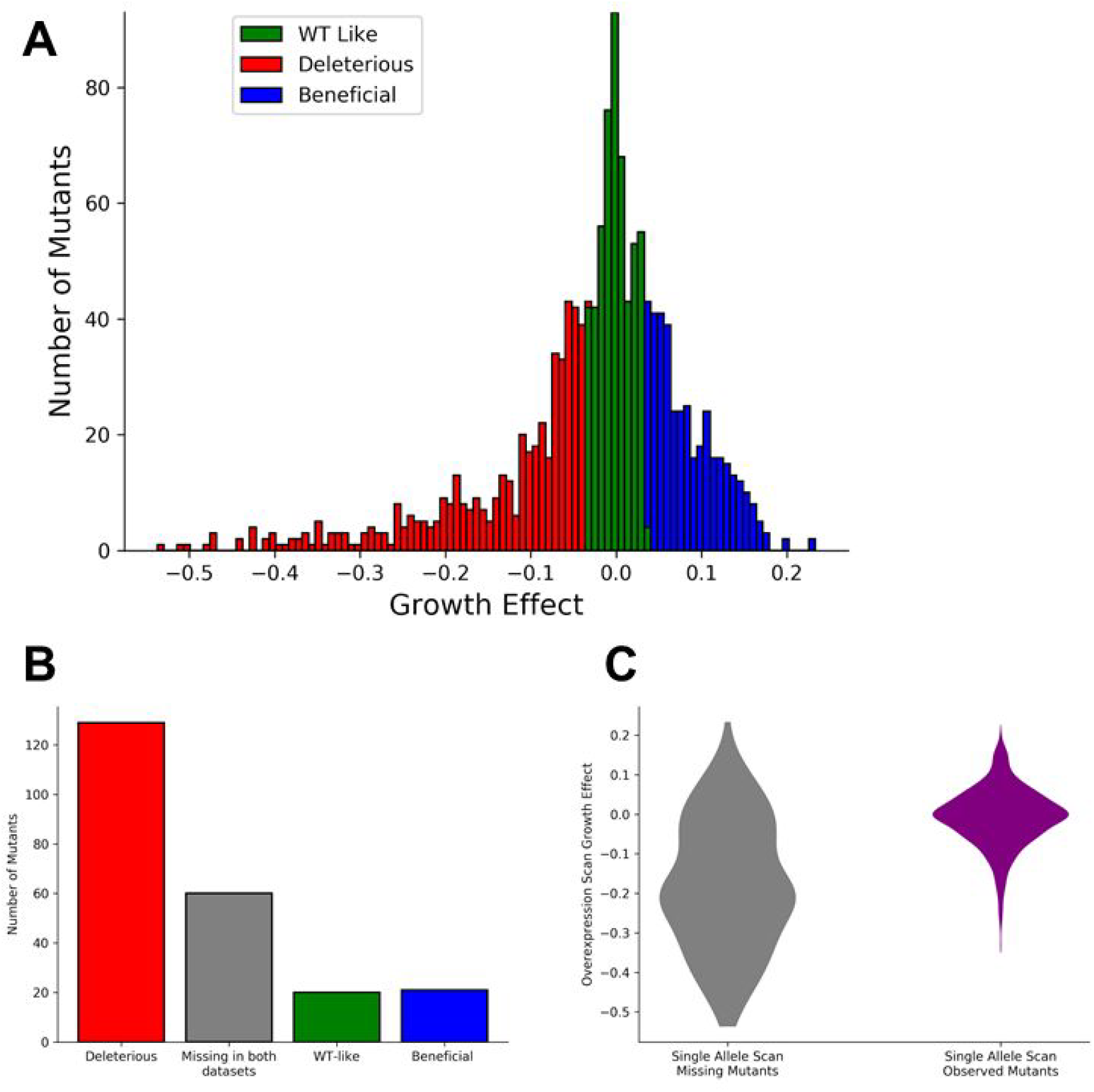
Over half of the alleles missing in the initial recessive scan can have deleterious effects when overexpressed in the presence of WT ubiquitin. A) The growth effects determined by the overexpression scan were binned into WT like alleles (green), deleterious alleles (red), and beneficial alleles (blue) by calculating a Z-score based on the distribution of WT synonymous alleles (Z > 4 and Z < −4). B) The bin of each allele missing in the single allele scan was determined.129 alleles fall into the deleterious bin, 60 alleles are missing in both datasets, 20 alleles are in the WT like bin, and 21 are in the beneficial bin. C) Mutants missing in the initial single allele scan (170 alleles) have a lower average fitness score in the overexpression scan than those observed in the single allele scan (1173 alleles). These two distributions are significantly different by a KS-test (p = 2.5*10^−44^)

Because these experiments were motivated by the observation that many mutants were depleted during the outgrowth of the recessive scans, we compared the overexpression populations to the population of missing alleles in the single allele scan (Figure 2B). Of the 230 alleles missing in the initial single allele scan, 129 alleles fall into the deleterious bin, 60 alleles are missing in both datasets, 20 alleles are in the WT like bin, and 21 are in the beneficial bin. Over half of the missing alleles in the single allele scan can be explained by their deleterious growth effects in the overexpression scan.

To further interrogate the relationship between mutants missing in the single allele scan and dominant growth effects, we compared the distribution of growth effects in the overexpression scan of two populations: those observed in the single allele scan and those unobserved in the single allele scan (Figure 2C). The distribution of growth effects for the 170 missing alleles is broad, with growth rates covering the whole range observed in the dominant scan. This distribution is enriched in negative scores, with a peak at ~-0.2. In contrast, the mutants observed in the single allele scan (1173 mutants) is tightly constrained with most mutants show no change in growth rate.

### Expression of WT Ub from both plasmids confirs a slight growth disadvantage

Upon examining the growth rates from the overexpression scan, we observed that mutations encoding stop codons grew faster than cells expressing two WT alleles. To quantify this effect, we compared the distribution of growth rates for WT synonymous mutants to that of stop codons (Figure 3A). These distributions are significantly different (KS-test, p = 3.1*10^−32^) with WT synonyms tightly constrained around a growth rate of 0 and stop codons more broadly distributed and with a peak at 0.11.

**Figure 3.**
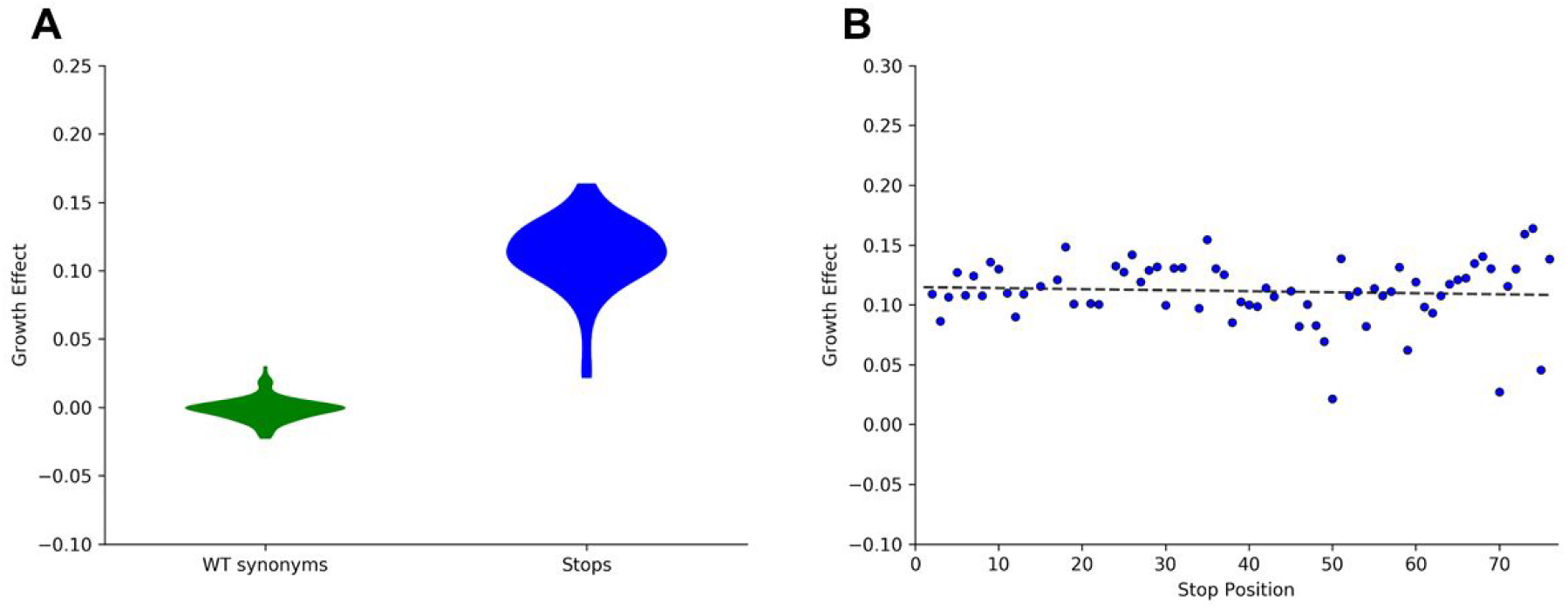
Stop codons appear beneficial in the overexpression scan due to overexpression of WT and functional alleles leading to the over-ubiquitination of the proteome. A) A violin plot showing the distributions of WT synonymous alleles (green) and stop codons (blue). These distributions are significantly different (KS-test, p = 3.1*10^−32^). B) The growth effect of overexpressing a mutant allele does not correlate with the protein length showing that this beneficial effect is not due to a reduced synthetic cost, but due to the non-functionality of the allele. (linear regression, Slope = −8.6*10^−5^, intercept = 0.11, R^2^ = 0.0052)

To address the hypotheses that this increase in growth rates of the stop codon population was due the synthetic cost of creating more ubiquitin molecules we plotted the mean stop codon fitness versus the position in the protein (Figure 3B). To our surprise, there is no change in growth rate as the position, and therefore the number of amino acids successfully translated, increases (linear regression, Slope = −8.6*10^−5^, intercept = 0.11, R^2^ = 0.0052).

These observations suggest that the increased growth rates of stop alleles is not due to a reduced protein synthetic cost but instead due to expressing additional WT ubiquitin leading to over ubiquitination of the proteome and a decrease in growth rate. This also explains the observation that other non-functional ubiquitin mutants, such as the terminal glycine mutants, appear to be beneficial in the overexpression scan.

### Strong growth effects in the overexpression scan appear throughout the ubiquitin structure

To examine the distribution of growth effects on the ubiquitin structure we colored each position in ubiquitin (1UBQ) based on the growth effect of a proline substitution. Proline was chosen because it both disrupts the protein fold as well as residue physio-chemistry. In contrast to the recessive scan, where strong deleterious growth effects were observed only on the face of ubiquitin containing the hydrophobic patch, we observed both strong deleterious and beneficial growth effects throughout the protein (Figure 4).

**Figure 4.**
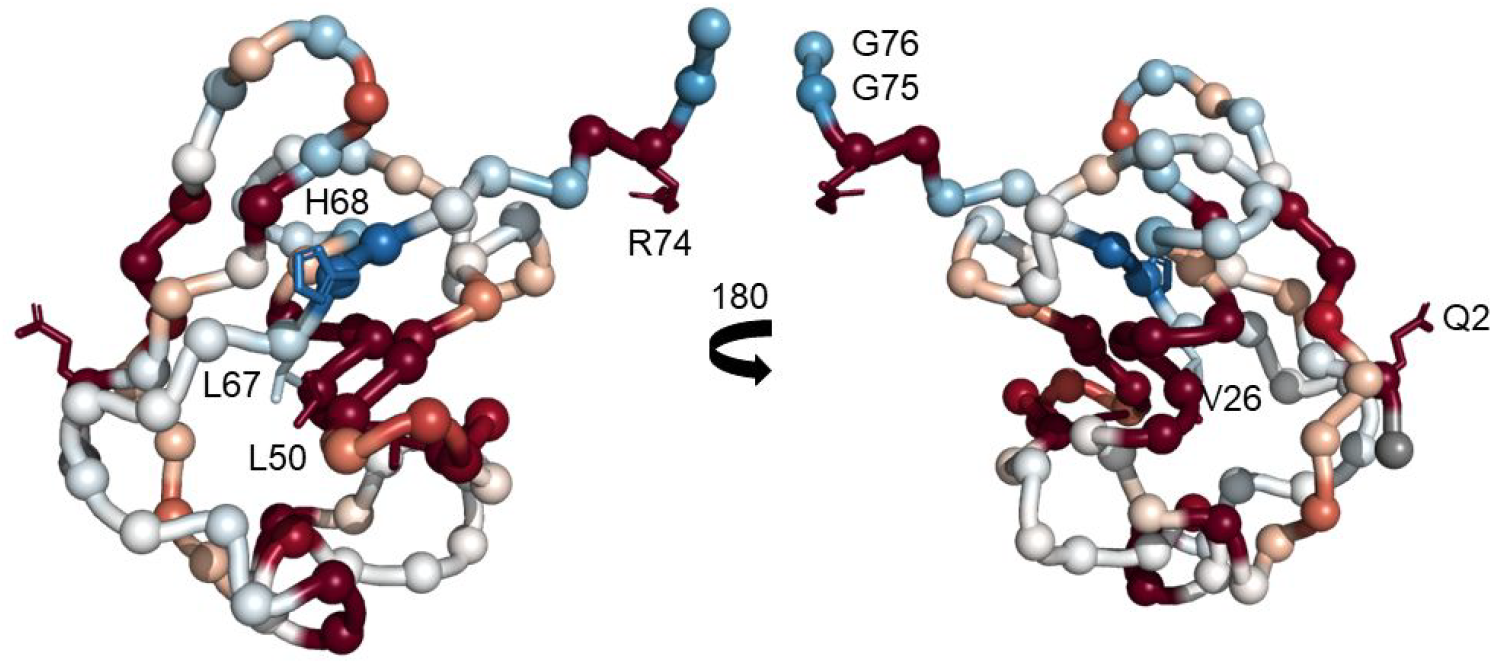
Positions exhibiting strong growth effects in the overexpression scan are distributed throughout the ubiquitin structure. The structure of ubiquitin (1UBQ) colored based on the growth effect of a proline substitution at that position with deleterious effects shown in red and beneficial effects shown in blue as in Figure 1. Positions with at least half of the observed substitutions showing a growth effect greater than 0.1 or less than −0.2 are highlighted (Q2, V26, L50, L67, H68, R74, G75, G76).

We focused on eight positions with large growth effects by highlighting those positions where at least half of the observed substitutions had a growth effect greater than 0.1 or less than −0.2. These positions include three core residues (V26, L50, L67), three previously identified sensitive face reissues (Q2, H68, R74), and the di-glycine motif (G75, G76). Intriguing, two of the core positions commonly show strong deleterious growth effects when mutated (V26, L50), suggesting that these mutations create functional but toxic proteins, while mutation at L67 leads to strong beneficial growth effects, suggesting that these mutations create non-functional proteins.

The strong beneficial effect of mutating the di-glycine motif fits with the observation that non-functional proteins appear as beneficial in the overexpression scan, as these residues are essential for E1 activation. The strong deleterious growth effect of mutating the adjacent R74 is intriguing. Like the di-glycine motif, R74 is inserted into the E1 active site during ubiquitin activation and is close to the E2 active site upon ubiquitin transfer. Previous data shows that mutants of R74 can be activated by E1(25) however the single allele growth effect scan contains few observations for mutations at position 74. The deleterious growth effects of mutants at this position may be due to the inhibition of ubiquitin transfer from E1 to E2 and E2 to E3 leading to a backup of the ubiquitin-proteasome system and dysregulated proteostasis.

## Discussion

This study represents the first use of DMS to study the growth effect of all possible point mutations when overexpressed in the presence of the WT allele. We observed growth rates that strongly corroborate our single allele scan, with the majority previously missing mutants showing deleterious growth effects in this overexpression scan. Additionally, the mean overexpression growth rate for mutants missing in the single allele scan is significantly lower than that of mutants observed in the single allele scan.

Interestingly it appears that non-functional alleles appear beneficial in the overexpression scan. This effect is most obviously seen in the distribution of stop codon growth effects, and is not due to a reduced synthetic load. Similarly, mutations of the di-glycine motif, that is essential for E1 activation, appear to be beneficial in the overexpression scan. Intriguingly this effect does not hold true for mutations at K48. Instead these mutations appear to have a mild deleterious effect. This suggests that these alleles are activated and ligated onto substrates, but because they cannot form K48 linked poly-ubiquitin chains, substrate proteins are not effectively turned over and proteostasis is perturbed even in the presence of WT ubiquitin.

Mapping these overexpression growth effects to the structure reveals that, unlike the single allele growth effects, there is not a sensitive and tolerant face of ubiquitin for dominant effects. Instead strong deleterious and beneficial mutations are distributed throughout the protein. We also identified eight positions with many large effects. While some are easily explained by basic ubiquitin biology or protein stability, mutations to R74 appear likely to inhibit the transfer of ubiquitin from E1 to E2 or E2 to E3 based on previous data and the position of the residue.

This study presents a novel approach to determine mutant allele growth effects in essential proteins based on overexpression in the context of the WT allele. Due to the ease of implementing this approach in parallel with a normal single allele DMS, we expect that this type of overexpression scan should become common.

## Materials and Methods

### Creation of the Ubiquitin Library

The saturating library of ubiquitin single point mutant alleles was generated using oligos containing NNN codons as previously described(12). Eight sub-libraries were created by mutating 10 amino acid long stretches in a KanMX4 marked high copy shuttle vector p427 with a Gal promoter. To facilitate sequencing during bulk competition, each variant of the library was tagged with a unique barcode via restriction digestion, ligation, and transformation into chemically competent E. coli.

### Association of mutant barcodes

Barcodes were added and associated with specific mutants as previously described(26). We performed paired-end sequencing of each sub-library using primers that read the N18 barcode as well as the ubiquitin coding sequence. To facilitate efficient Illumina sequencing, we generated PCR products that were less than 1kb in length for sequencing. The library plasmids were harvested by miniprep (Zymo Research) and the target sequence amplified with 11 cycles of PCR to generate products for Illumina sequencing. The resulting PCR products were sequenced using an Illumina MiSeq. Low-quality reads (Phred scores <10) were removed and for each barcode that was read more than three times we generated a consensus of the ubiquitin mutant sequence.

### Bulk Competition

Equal molar quantities of each sub-library were mixed to form a pool of DNA containing the entire ubiquitin library. The plasmid library was transformed into W303 using the lithium acetate method. Transformations were performed until enough transformants were generated to give each barcode an average of 10X coverage in the library. Transformants were recovered for 12 hours in SD media and then outgrown in log phase for 48 hours in SD+G418

To start the bulk competition, cells in log growth were washed with S-Raf+G418 and then grown in S-Raf+G418 for 16 hours until they reached mid-log phase. A sample was collected to serve as the zero time point of the selection. Cells were then diluted into S-RafGal+G418 and grown in competition for 24 hours. Samples were taken after 2 hours, 5 hours, 8 hours, 11 hours, 14 hours, 17 hours, 20 hours, and 24 hours. The culture was maintained in log phase with a population size of at least 10^9^ throughout the competition.

### DNA Preparation and Sequencing

Plasmid DNA from each sample was isolated via lysis and mini prep (Zymo Research). Three PCR steps were performed to amplify the DNA for sequencing using Phusion polymerase (NEB). First primers targeting the p427Gal vector were used to amplify DNA containing the N18 barcode sequences. These products were gel purified (Zymo Research) and used as templates for the second PCR with a primer containing the MmeI restriction site. This second product was purified by silica column (Zymo Research) and digested with MmeI (NEB). The digested product was gel purified and then adaptor sequences containing the sample identifier sequences were ligated to the product. This product was gel purified and a final PCR was performed to add the p5, p7, and PE1 sequences necessary for Illumnina sequencing. These final PCR products were purified two times over silica columns (Zymo Research) and quantified using the KAPA SYBR FAST qPCR Master Mix (Kapa Biosystems) on a Bio-Rad CFX machine. Samples were pooled and sequenced on an Illumina NextSeq in single-end 100 bp mode.

### Analysis of Bulk Competition Sequencing Data

Illumina sequence reads were filtered for Phred scores >20 and strict matching of the sequence to the expected template and identifier sequence. Reads that passed these filters were parsed into time points based on the identifier sequence. For each time point identifier, each unique N18 barcode read was counted and the frequency of each allele in each time point calculated. A cumulative count for each amino acid variant was generated by summing the counts of all N18 barcodes that mapped to that amino acid variant.

### Determination of Selection Coefficient

The frequency of each variant in the library determined at each time point. Selection coefficients were calculated by finding the slope of the linear fit between generation time and the log of allele frequency. These selection coefficients were then normalized to the mean all WT synonymous alleles (s=0).

## Acknowledgements

This work was supported by grants from the National Institutes of Health (R01-GM112844 to D.N.A.B. and F32-GM119205 to D.M. T32-CA130807-10)

